# Regulated extracellular matrix trafficking shapes cell growth during cartilage morphogenesis

**DOI:** 10.1101/2022.06.08.495337

**Authors:** Daniel S. Levic, Gokhan Unlu, David B. Melville, Ela W. Knapik

**Author notes:** Correspondence: Ela W. Knapik, 1165 Light Hall, 2215 Garland Ave., Nashville, TN 37232.

## Abstract

Craniofacial malformations are present in more than one third of all congenital syndromes, but the pathogenesis of skeletal dysmorphology is poorly understood. Here, using an unbiased forward genetics approach in zebrafish, we identified a mutation in *erc1b* that leads to craniofacial defects, including micrognathia and hypertelorism caused by impaired cartilage and bone growth. To date, *ERC1* has not been considered a candidate gene for craniofacial syndromes. Using live *in vivo* imaging, genetic depletion and replacement experiments, and transgenic approaches, we interrogated *erc1b* function. We found that Erc1b regulates extracellular matrix (ECM) trafficking required for the highly conserved “stack of coins” organization of chondrocytes in cartilage that is essential for skeletal growth and integrity. Erc1b functions cell-autonomously at the chondrocyte cell cortex to regulate traffic of ECM and plasma membrane expansion in a microtubule dependent manner during isometric cell growth. Disruption of Erc1-Rab8-Kinesin-1 axis leads to failure of cartilage maturation, endochondral bone formation and ultimately chondrocyte cell death. Our study identifies Erc1b as a candidate genetic factor for craniofacial syndromes.

## Introduction

Craniofacial morphogenesis is a complex series of developmental processes that is afflicted in many human congenital disorders. Morphological defects of the jaw and skull are highly prevalent in human syndromes^1^, likely because multiple cell lineages and gene regulatory networks contribute to the patterning, formation, and growth of the facial skeleton^2^. During embryonic development, cartilage is a critical precursor for parts of the facial skeleton, providing structural integrity and functioning as a template and scaffold for bony tissue that will later define craniofacial architecture. Given the complex genetic landscape of human craniofacial syndromes^3,4^, detailed animal model studies of facial cartilage development are needed to better understand human craniofacial pathologies^5^.

Studies in chick, mice, and zebrafish have helped to shed light on mechanisms of craniofacial cartilage development^2,5^. Cranial neural crest cells (NCCs) populate pharyngeal arches^6^ where they condense to form pre-cartilage intermediates that define the future morphologies of individual skeletal elements^7,8^. After NCCs differentiate into chondrocytes, newly formed cartilage undergoes rapid growth that drives jaw extension^7^. This rapid growth phase of cartilage development is driven largely by cartilage ECM secretion^9–11^ and convergent extension as chondrocytes rearrange from small, round cell clusters into organized stacks of elongated cells^7^. Chondrocyte stacking, which was first documented nearly a century ago^12^, is critical for craniofacial and axial cartilage growth in humans^13^, mice^14,15^, and zebrafish^16–19^. However, the fundamental question of which pathways contribute to cartilage architecture and chondrocyte shape changes largely has not been examined.

Here, we explore the roles of ECM trafficking in chondrocyte during cell growth and cartilage morphogenesis. We find that in the absence of the cortical scaffolding protein Erc1b, or upon mosaic inhibition of Rab8 or Kinesin-1, chondrocytes progressively decrease in cell width, which is suppressed when Rab11-dependent recycling is inhibited. Cell shape dysregulation impairs craniofacial cartilage growth and later induces chondrocyte cell death, further compromising cartilage integrity and leading to endochondral bone defects. Loss of Erc1-dependent exocytic trafficking does not impair chondrocyte convergent extension, suggesting that cell shape dynamics and morphogenetic cell rearrangements are regulated independently during cartilage growth. We propose that chondrocyte cell growth is shaped by a membrane recycling pathway during a phase of rapid biosynthetic cargo delivery to the plasma membrane in developing cartilage.

## Results

### The *kimble* mutant as a genetic model for craniofacial morphogenesis

To investigate cellular mechanisms underlying craniofacial growth and morphogenesis, we utilized a chemically-induced zebrafish mutant, *kimble* (*kim*^*m533*^)^20^, which exhibits a specific defect in jaw morphology at post-embryonic stages (5 dpf, Fig. 1A-D). The *kimble* mutant phenotype mostly affects the craniofacial skeleton, while the axial skeleton and body stature is similar in mutants and wild-type (WT) siblings (Fig. 1A). Moreover, the earliest stages of craniofacial development, including neural crest cell (NCC) migration, pharyngeal arch patterning, and chondrocyte cell specification, are unaffected in *kimble* mutants at 3 dpf (data not shown; Fig. 1E; Supp. Fig. 1A,B). Gross defects in mutant craniofacial morphology were evident in Alcian blue stained skeletal preparations at 4 dpf, with *kimble* mutants exhibiting cartilage degradation apparent in the otic capsule and ethmoid plate (Fig. 1G, arrow and data not shown). Loss of cartilage in *kimble* mutants was likely a consequence of substantial chondrocyte cell death (Fig. 1H,I) since chondrocyte specification (Fig. 1E) and cartilage formation at 3 dpf (Supp. Fig. 1A,B) were unaffected. Defects in craniofacial cartilage integrity led to impaired bone formation at later stages, with cartilage-derived bones absent (hyomandibular bone, HMB) (Fig. 1J), while intramembranous bones were unaffected (opercle, OP) (Fig. 1J). In summary, *kimble* mutants present broad craniofacial dysmorphologies, including micrognathia and hypotelorism, stemming from cartilage and bone defects that occur after pharyngeal arch patterning and chondrocyte specification.

**Figure 1.**
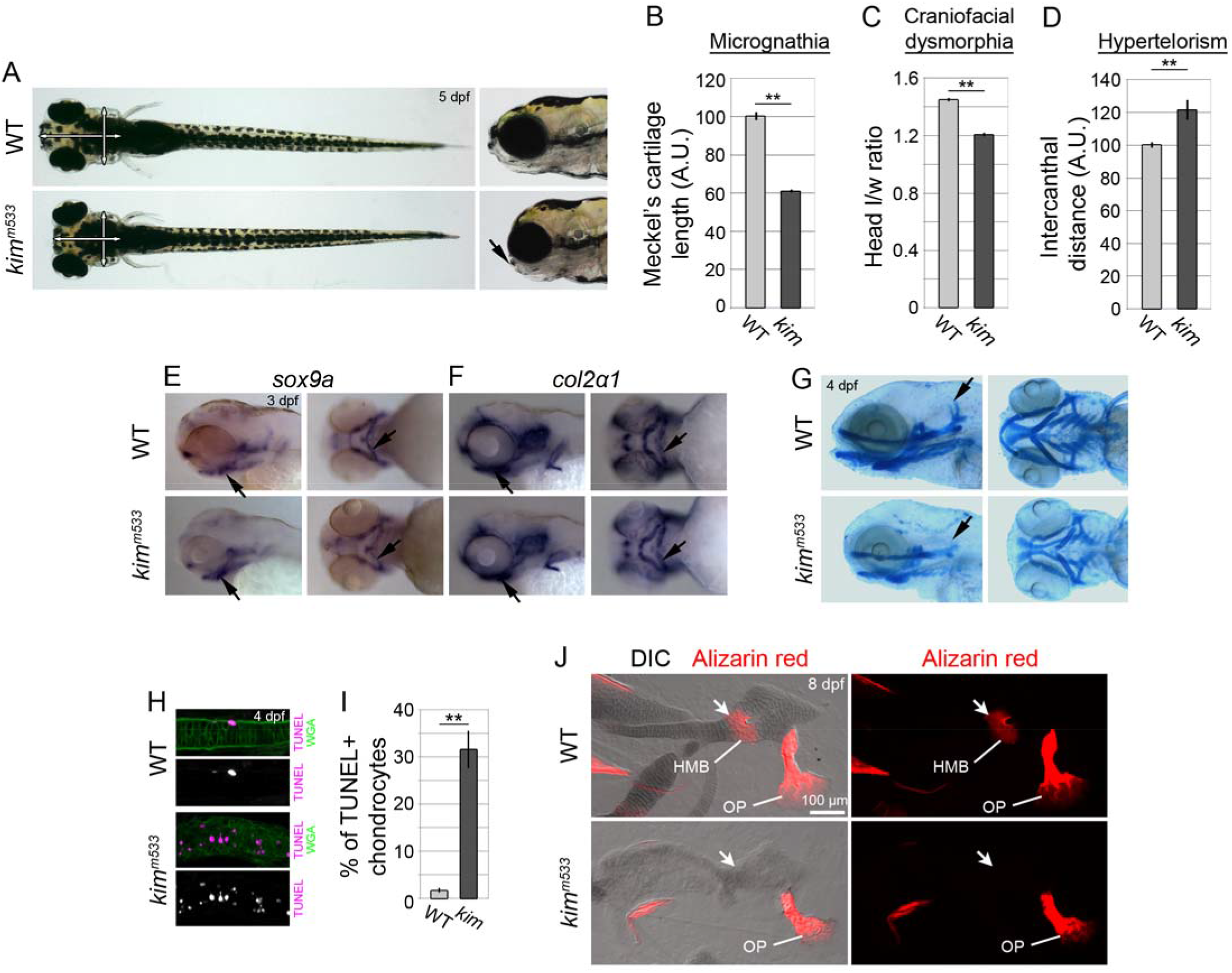
*kimble*^*m533*^ mutants present broad craniofacial skeletal deficits after chondrocyte differentiation. (A) Live images of WT sibling and kimble (*kim*^*m533*^) mutant 5 dpf larvae. Live *kimble* mutants are indistinguishable from WT siblings at early stages but exhibit craniofacial malformations at post-embryonic stages (5 dpf, arrowhead). White arrows denote areas used for quantitative measurements. (B-D) Quantitative analysis of craniofacial morphogenesis. Micrognathia (B) was assessed using Alcian blue-stained skeletal preparations (G), and craniofacial dysmorphia (C) and hypertelorism (D) were analyzed on live larvae (A). n ≥ 3 animals(E-F) Chondrocyte differentiation and patterning is unaffected during embryogenesis in *kimble* mutant embryos. *sox9a and col2a1* mRNA expression (arrows) were analyzed using whole mount in situ hybridization. (G) *kimble* mutants exhibit skeletal deformation and degeneration at the early post-embryonic stage (4 dpf, otic capsule cartilage, arrows). (H-I) *kimble* mutants exhibit widespread chondrocyte cell death during cartilage degeneration (4 dpf) as assessed by TUNEL. n > 100 cells from at least 5 animals. (J) *kimble* mutants exhibit a specific defect in cartilage-derived bones at late larval stages (hyomandibular bone, HMB, arrows), not intramembranous bones (opercle, OP). 8 dpf larvae were stained with Alizarin red, and skeletal preparations were dissected, flat-mounted, and imaged with DIC and fluorescence microscopy. **p<0.01

### The *kimble*, mutation ablates erc1b function

To understand how the *kimble* mutation impacts cartilage growth and morphogenesis at a molecular level, we used a positional cloning strategy to identify the mutation site^21^. After restricting the critical region to a 500 kb interval containing a single gene on the proximal arm of chromosome 4 (Fig. 2A), we identified a C>T transition at base pair 757 of *erc1b* in *kim*^*m533*^ mutants, resulting in a Q253X nonsense mutation that is predicted to truncate nearly 75% of the highly conserved protein (Fig. 2B). Double-blinded genetic replacement experiments via mRNA injection confirmed that *erc1b* expression was sufficient to restore normal jaw morphogenesis in *kimble* mutants (Fig. 2C), and mosaic transgenic replacement assays demonstrated that *erc1b* regulates chondrocyte survival cell-autonomously (Fig. 2D-F). Consistent with cell autonomous function, whole mount *in situ* hybridization using riboprobes against the *erc1b* 3’ UTR showed enriched expression in developing cartilage, with prominent expression in the pharyngeal arches (Fig. 2G, arrows). *Erc1b* mRNA expression in chondrocytes was confirmed by RNA-Seq of fluorescent activated cell sorted (FACS) head chondrocytes (Unlu et al., unpublished observations). Transcripts of *erc1b* (ENSDARG00000076014), which correspond to the *kimble* gene on chromosome 4, were 1.9-fold more highly expressed in head chondrocytes relative to non-skeletal tissue. Moreover, the related paralog *erc1a* (ENSDARG00000009941) on chromosome 25, was not highly expressed in head chondrocytes, with *erc1b* 20.4-fold more highly expressed in chondrocytes than *erc1a*, which was instead highly enriched in non-skeletal tissues (86-fold enriched). These tissue-specific expression differences among the zebrafish *erc1* paralogs are likely the reason that *kimble* mutants exhibit morphological defects mostly specific to the craniofacial skeleton.

**Figure 2.**
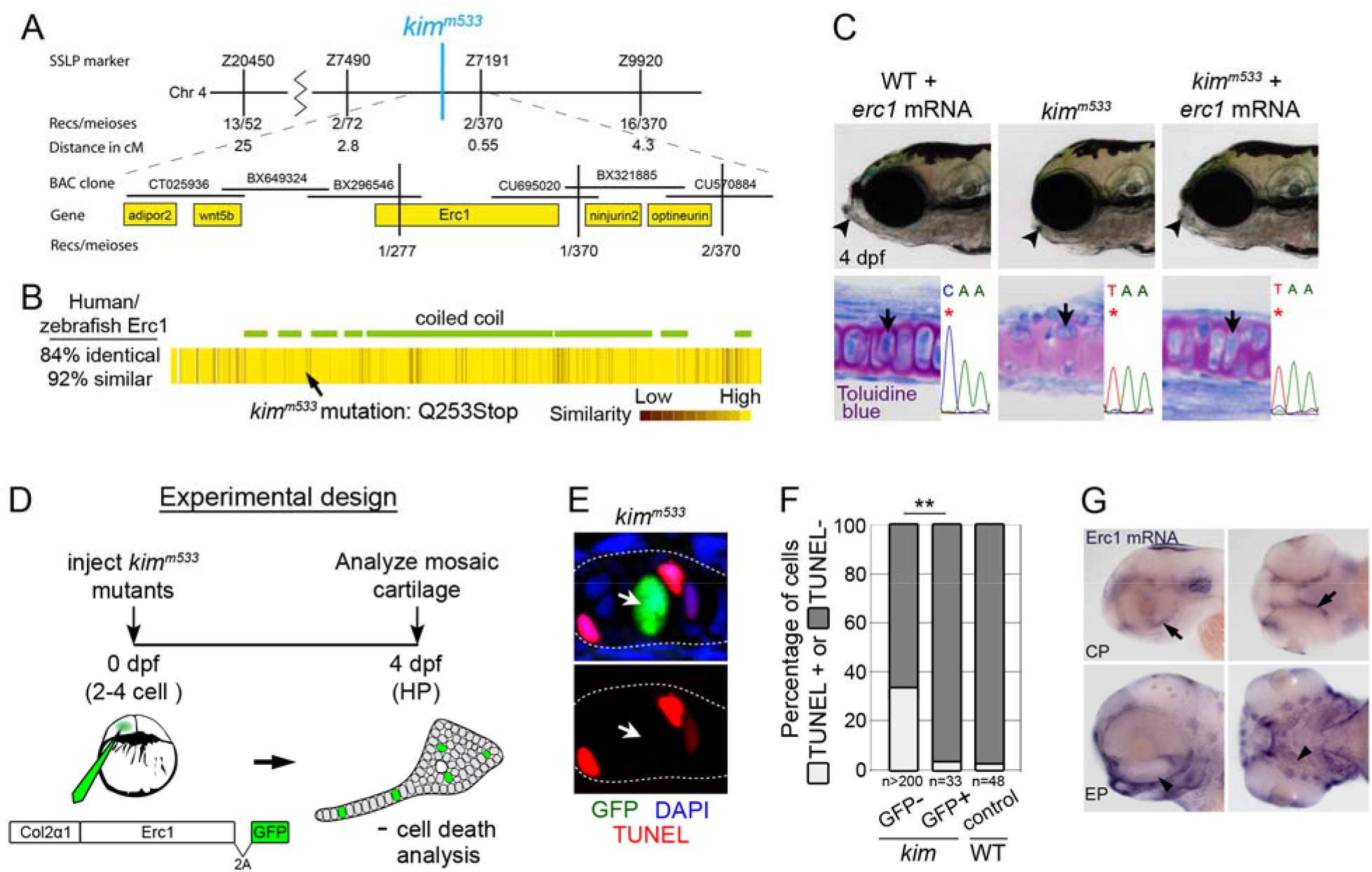
Loss of *erc1b* underlies craniofacial morphogenesis and chondrocyte cell survival defects in *kimble* mutants. (A) Schematic illustration of the *kimble* mutation critical interval, which contained only *erc1*. (B) Heat map schematic of Erc1 amino acid conservation was generated using ClustalW2 and Jalview. (C) Expression of WT *erc1b* mRNA in *kimble* mutant embryos rescues jaw morphogenesis (arrowhead) and chondrocyte cell shape (arrow). Analyzed embryos were genotyped by sequencing (asterisks mark the SNP). (D) Schematic illustrating the experimental design for mosaic rescue assay used for testing Erc1b cell-autonomous function. Rescue construct *col2*α*1:erc1-v2a-eGFP* drives expression of untagged Erc1b under the collagen 2α1 promoter in chondrocytes, and 2a-eGFP reporter marks transgenic cells. (E-F) *erc1b* is required cell-autonomously chondrocyte cell survival in *kimble* mutants. n values are number of cells analyzed from 9 animals. (G) Endogenous *erc1b* mRNA expression is enriched in developing cartilage, including the neurocranium (arrow) at 2.5 dpf (CP) and pharyngeal skeleton (arrowhead) at 3 dpf (EP). **p<0.01

### Erc1b regulates chondrocyte cell shape changes during a phase of rapid cell growth and ECM secretion

As the only cell type within cartilage, chondrocytes impact craniofacial growth not only through cartilage ECM secretion but also by morphogenetic behaviors such as convergent extension via cellular intercalation^7^. Thus, because gross craniofacial defects were not apparent immediately after chondrocyte specification in *kimble* mutants (Supp. Fig. 1A,B), we asked whether severe dysmorphology and cartilage degradation could be a consequence of altered chondrocyte organization at earlier stages. To address this question, we analyzed intercalating chondrocytes during and immediately after differentiation using the simple rod-shaped symplectic (SY) cartilage as a model. SY cartilage forms as clusters of condensed, non-proliferating cells rearrange into stereotypic stacks of elongated chondrocytes^7^.

We first analyzed SY cartilage in wild types and divided its formation into three phases, defining a developmental staging system for our study. First, at the condensation phase (CP, 2.5 dpf), WT chondrocytes were arranged in clusters of small, round cells that were Alcian blue-negative before the onset of ECM production (Fig. 3A). In the subsequent elongation phase (EP, 3 dpf), WT chondrocytes were arranged into organized stacks and had undergone cell elongation, as shown by increased cell length-to-width (l/w) ratios (Fig. 3B,G,H). EP chondrocytes were also Alcian blue-positive, indicating that ECM proteins are synthesized and secreted during this phase. Finally, at the isometric growth phase (IGP, 3.5 dpf), chondrocytes exhibited additional cell growth but without increases in l/w ratios, and they were also separated by additional ECM compared to earlier phases (Fig. 3C,G,H).

**Figure 3.**
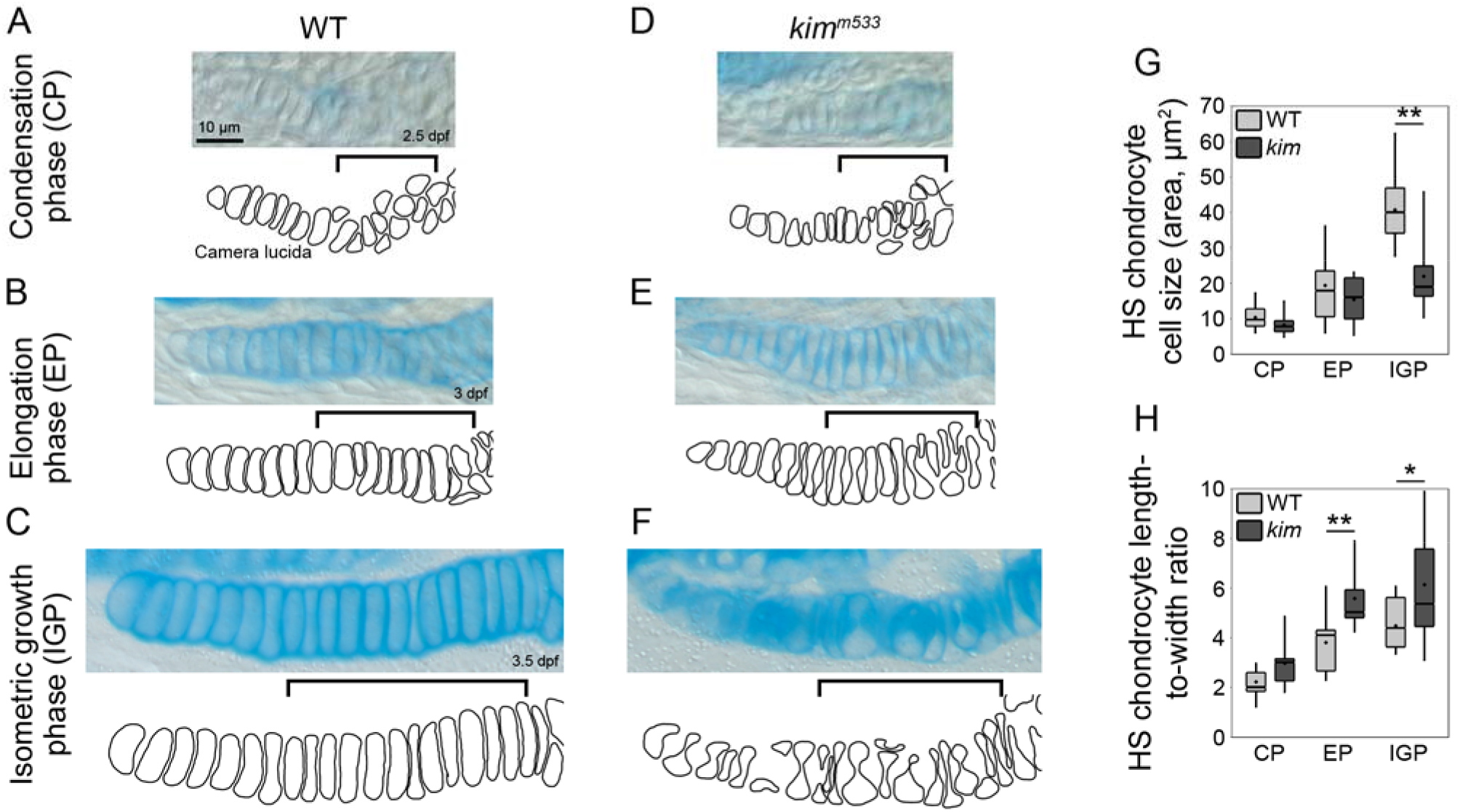
Erc1b deficient chondrocytes dysregulate cell shape changes after chondrocyte intercalation. (A-F) *kimble* mutants exhibit cell shape and organization defects immediately after chondrocyte differentiation at the onset of cartilage ECM secretion. DIC images and camera lucida cell outlines of Alcian-blue stained hyosymplectic (HS) WT cartilage were used for quantitative analysis. Brackets indicate chondrocytes that resolve from progenitor clusters to stacked chondrocytes during differentiation. The CP-EP transition in wild types is characterized by cell elongation and extracellular matrix (ECM) secretion (A,B), while the EP-IGP transition is accompanied by isometric cell growth (B,C). CP-EP *kimble* chondrocytes stack (intercalate) but are disproportionally elongated due to constriction along the lateral cell edge (D,E). EP-IGP *kimble* chondrocytes grow at polarized ends, resulting in an hourglass cell shape (F). (G,H) Quantification of cell size and shape changes during differentiation. n = 13 chondrocytes. *p<0.05, **p<0.01

In *kimble* mutants, CP chondrocytes were indistinguishable from wild types (Fig. 3D,G,H), and EP chondrocytes rearranged into stacks concurrent with ECM secretion (Fig. 3E). However, EP *kimble* chondrocytes exhibited striking cell shape defects, forming elongated but highly constricted cells that had much greater l/w ratios than wild types (Fig. 3E,G,H). These cell shape defects increased in severity from the EP-IGP transition in *kimble* mutants, and polarized cell ends appeared expanded in width relative to earlier stages, while the lateral sides remained highly constricted (Fig. 3F-H). We confirmed this complex cell shape defects using confocal microscopy and 3D cell surface rendering (Fig. 4A,B, Supp. Videos 1-2). *kimble* chondrocytes appeared to grow circumferentially at polarized ends but were substantially decreased in cell width. To test whether cell shape defects impact cartilage growth at this early stage, we imaged hyosymplectic (HS) cartilage in embryos using confocal imaging of wheat germ agglutinin (WGA) stained embryos (Fig. 4C,D). We then performed 3D surface rendering of whole HS cartilage to measure organ volume, which showed that *kimble* cartilage was 34.4% smaller than wild types (Fig. 4C-E, Supp. Videos 3-4). These data indicate that cartilage morphogenesis depends on properly executed cell shape changes during a phase of rapid cell growth and ECM secretion (Fig. 3A-H, Fig. 4F). Thus, although *kimble* mutants exhibit normal NCC migration, pharyngeal arch patterning, chondrocyte differentiation, and convergent extension, the compromised cell shape changes significantly impair cartilage growth.

**Figure 4.**
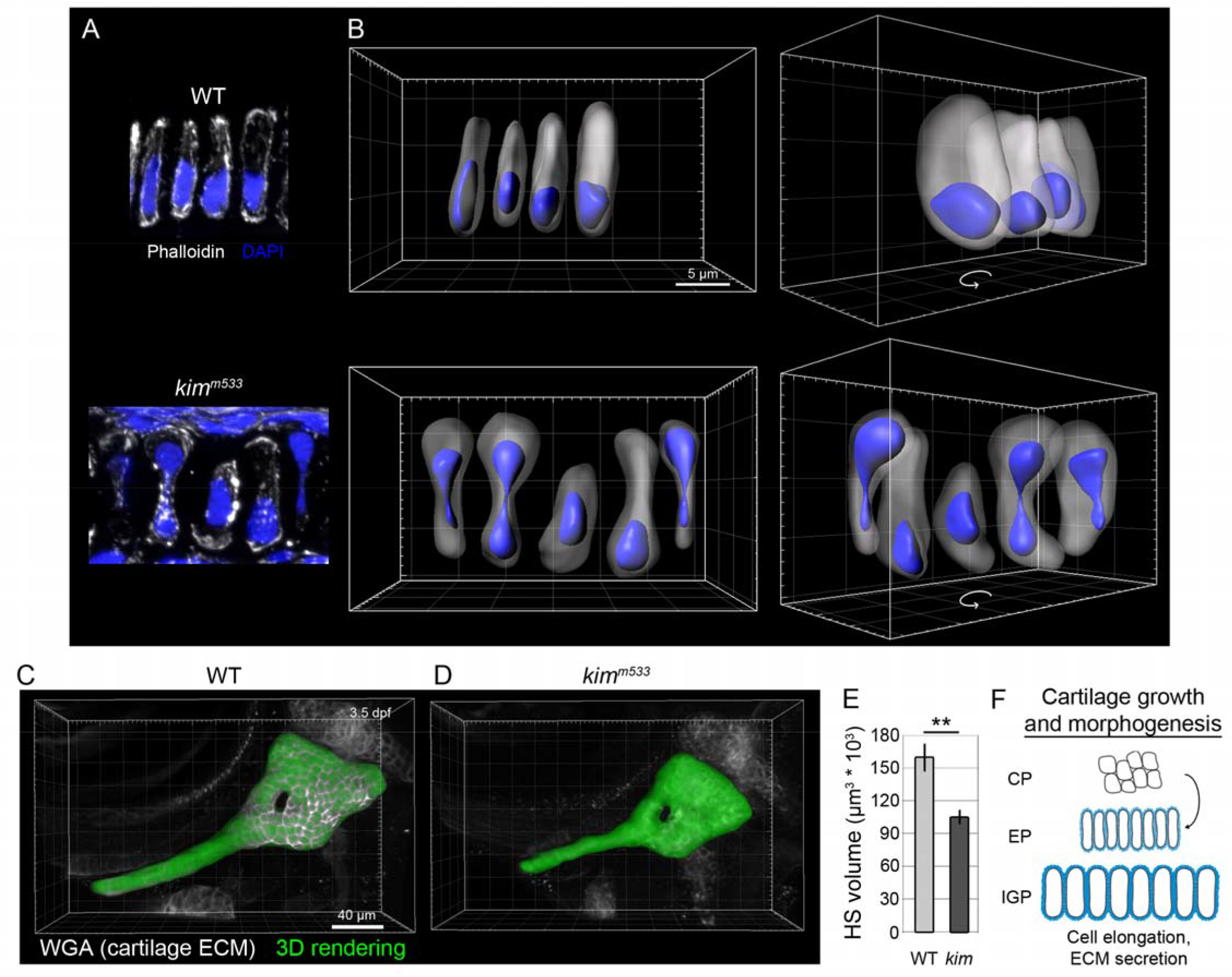
Chondrocyte cell shape dysregulation impairs cartilage growth and morphogenesis. (A) Fluorescence microscopy and (B) 3D rendering of cell and nuclear shape of stacked chondrocytes. WT chondrocytes exhibit a characteristic coin-shape and are arranged in a stacked organization, while *kimble* chondrocytes grow circumferentially at polarized ends, resulting in a complex hourglass cell shape with the nucleus often constricted in the midline of the cell. EP stage (3 dpf) embryos were stained with phalloidin and DAPI, imaged by structured illumination microscopy, and analyzed with 3D rendering using Imaris. (C-E) 3D rendering and volume measurements of hyosymplectic (HS) cartilage at 3.5 dpf (IGP). Confocal z-stacks of wheat-germ agglutinin (WGA) stained embryos were used for rendering and quantification. n ≥ 4 animals. (F) Schematic illustrating cell behavior changes accompanying chondrocyte differentiation. Cell shape changes and ECM secretion are temporally linked during chondrocyte cell growth and cartilage morphogenesis. **p<0.01

### Loss of cell shape integrity impairs chondrocyte cell survival

Our data show that loss of *erc1b* leads to severe cell shape defects after chondrocyte differentiation (EP, 3 dpf, Fig. 3E), impairing cartilage growth (IGP, 3.5 dpf, Fig. 4E) and later leading to widespread chondrocyte cell death (4 dpf, Fig. 1H,I), which can be rescued cell-autonomously (Fig. 2D-F). Therefore, we asked whether chondrocyte cell death is a direct consequence of cell shape dysregulation. To test this possibility, we attempted to manipulate aberrant cell shape changes in *kimble* mutants by destabilizing cytoskeletal components, which have been shown to play an active role in cell shape changes in many cell-types^22–24^. Since we found that disrupting cortical f-actin (Supp. Fig. 2A,B) did not alter chondrocyte cell shape in either *kimble* mutants or wild-type siblings (Supp. Fig. 2C,D), we focused on microtubules. We treated embryos with nocodazole from 2.5-3.5 dpf (CP-IGP, Fig. 5A), using a low concentration (0.6 µM) that exhibited minimal toxicity after 24 hrs treatment. While this regimen only partially destabilized microtubules (Fig. 5B), *kimble* chondrocyte cell shape defects were mostly suppressed and mutant cells exhibited length-to-width ratios comparable to wild types (Fig. 5C,D). However, while cell constriction in *kimble* mutants was suppressed by microtubule destabilization, paclitaxel (20 µM) mediated microtubule stabilization in wild types did not recapitulate *kimble* mutant cell shape defects (Fig. 5C,D). Thus, microtubule stability or dynamics alone are likely not the cause of chondrocyte cell shape dysregulation in *kimble* mutants. We next tested whether suppressing cell shape defects in *kimble* mutants rescued cell death. Our results showed that partially destabilizing microtubules led to more than an 80% reduction in TUNEL+ *kimble* chondrocytes compared to DMSO-treated controls (Fig. 5E,F). These data indicate that cell shape regulation after chondrocyte differentiation is critical for maintaining proper cell survival and promoting cartilage growth.

**Figure 5.**
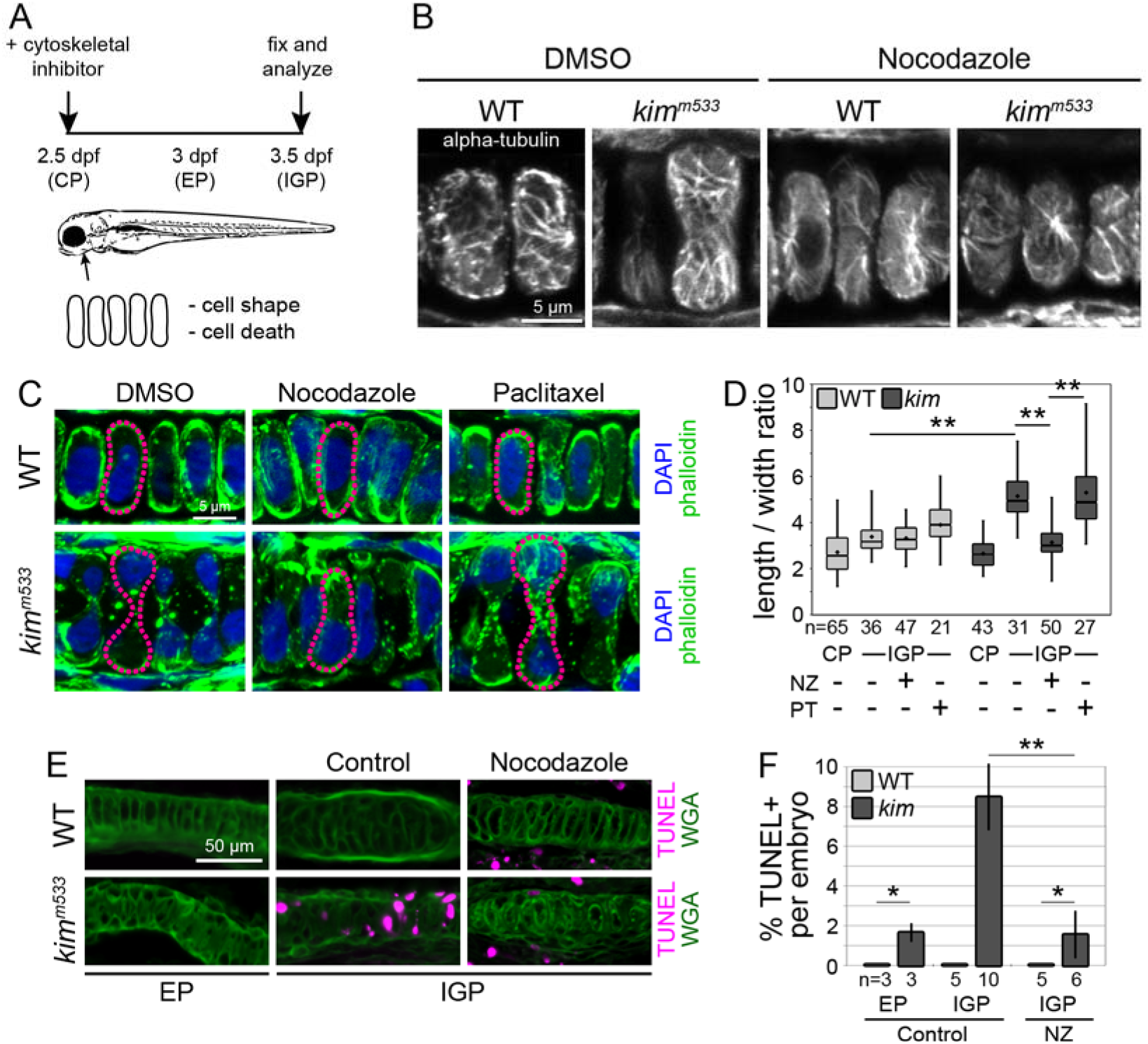
Impaired microtubule-dependent cell shape regulation leads to chondrocyte cell death. (A) Experimental design for testing the role of cytoskeletal components in chondrocyte cell shape regulation. (B) Imaging of microtubules (alpha-tubulin) after control treatment (DMSO) or microtubule destabilization (nocodazole, NZ). Microtubules are partially destabilized after 24 hr treatment with low concentration of NZ (0.6 µM). (C) Imaging of cell (phalloidin) and nuclear (DAPI) shape after treatment with 1% DMSO (control), 0.6 µM NZ, or 20 µM paclitaxel (PT) from CP-IGP. Destabilizing microtubules (NZ) rescues *kimble* chondrocyte cell shape defects, while stabilizing microtubules (PT) in WT embryos does not phenocopy *kimble* cell shape. Magenta outlines indicate representative chondrocytes. (D) Quantification of cell shape after indicated treatment. n values are number of cells analyzed from at least 3 animals. (E,F) TUNEL assay and quantification of chondrocyte cell death during chondrocyte differentiation and growth. Destabilizing microtubules (NZ treatment) rescues *kimble* chondrocyte cell death (E) as well as shape (C). n values are number of animals with at least 100 chondrocytes analyzed. *p<0.05, **p<0.01

### Erc1b regulates exocytic vesicle trafficking at the chondrocyte cell cortex

To investigate how Erc1b influences chondrocyte function and cartilage growth we sought to determine its subcellular localization in craniofacial chondrocytes. Erc1b localization has been shown to vary within different cultured cell lines ^25–28^, but its distribution in chondrocytes has not been reported. We obtained a commercially available monoclonal antibody against a human ERC1 peptide that is perfectly conserved with the zebrafish protein, and as a control we first showed that the antibody recognizes zebrafish Erc1b using western blot detection of zebrafish lysates overexpressing a GFP-Erc1b fusion construct (Fig. 6A, Supp. Fig. 3A). We also performed western blot analysis of endogenous Erc1b in *kimble* mutant lysates and observed loss of one of two closely migrating bands at approximately 130 kDa (Fig. 6B; Supp. Fig. 3B), consistent with predicted molecular weight. The remaining bands in *kimble* lysates likely represent Erc1a and Erc2, which are not expressed in chondrocytes since *kimble* cartilage shows no immunoreactivity by IF (Fig. 6C) and their transcripts were not significantly detected in chondrocytes by RNA-seq (our unpublished observations). We next performed IF and confocal imaging to determine Erc1b subcellular localization. Our results showed that endogenous Erc1b is present at the cell cortex in discrete puncta within WT chondrocytes (Fig. 6D, arrows). Subsequent 3D rendering (Imaris) of whole cartilage confocal z-stacks followed by distance mapping of Erc1b voxels relative to the plasma membrane (PM, caax-eGFP) showed that most Erc1b puncta were present at the cortical region (93.8 ± 4.7% were ≤ 1 µm from the PM, 95% CI, n=145) (Fig. 6E,F, Supp. Video 5). In summary, our data show that Erc1b localizes to the chondrocyte cell cortex and functions cell-autonomously to regulate shape changes during rapid cell growth phase.

**Figure 6.**
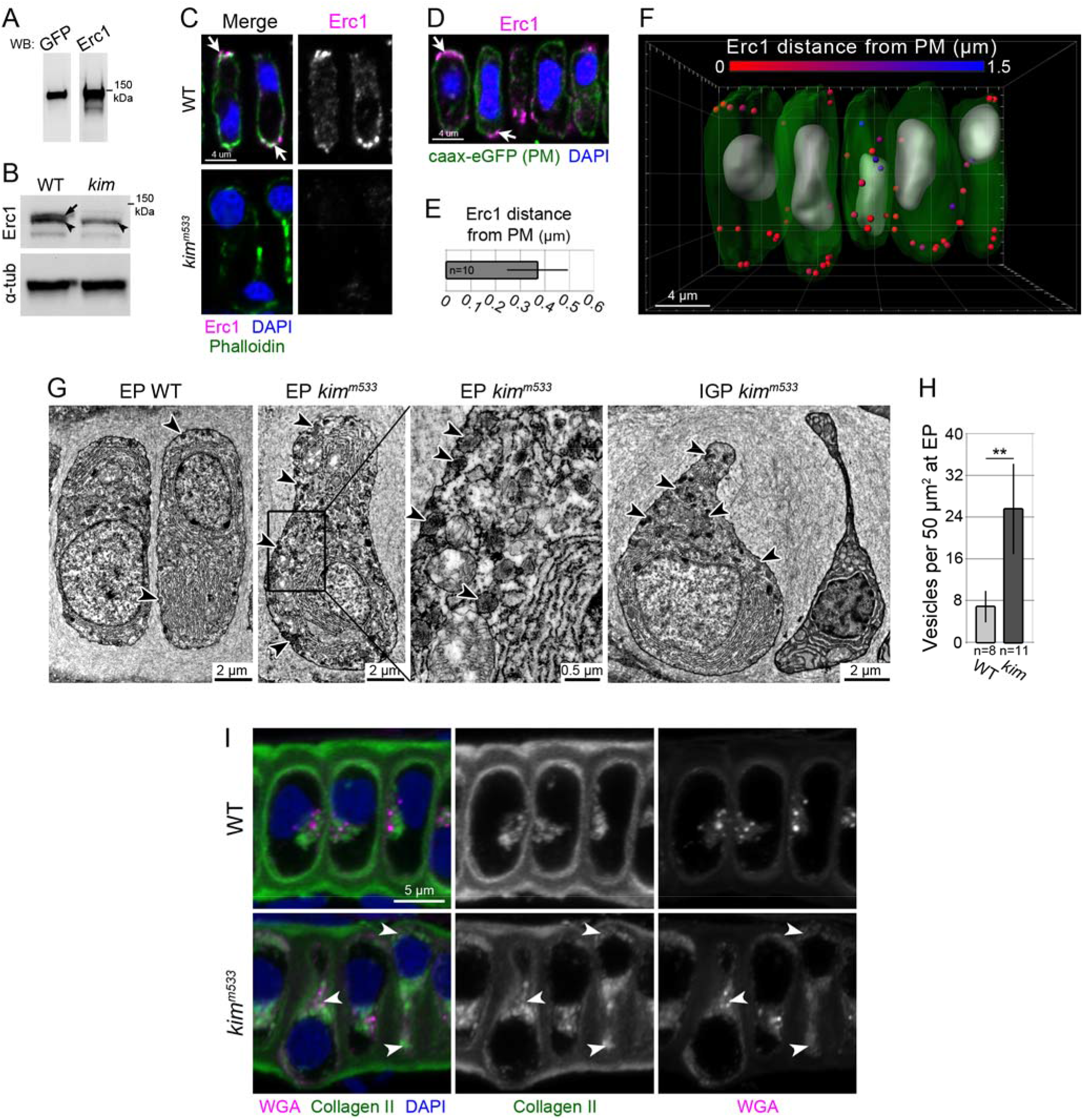
Regulation of cortical cargo trafficking is required for chondrocyte cell shape regulation and cartilage morphogenesis. (A) Erc1 antibody recognizes the zebrafish fusion protein in western blots of *eGFP-erc1b* injected embryo lysates. (B) Western blot analysis of endogenous Erc1 protein expression in WT and *kimble* lysates. *kimble* lysates show loss of immuno-reactivity of one of the two prominent bands (arrow) close in molecular weight between 100-150 kDa. The second band (arrowhead) is unaffected. Uncropped blot images are in Supp. Fig. 3A,B. (A,B). (C) Erc1b protein expression is lost in *kimble* chondrocytes. Cryosections of 3 dpf embryos were imaged with confocal microscopy using identical settings and processing. WT image is from a sibling of the *kimble* mutant. (D) Endogenous Erc1 protein (magenta) is localized in close proximity to the chondrocyte plasma membrane (PM, green, marked by *col2*α*1:caax-eGFP*). (E,F) Quantification and 3D reconstruction of Erc1b localization using Imaris. Spheres representing Erc1 voxels are pseudo-colored according to their distance from the PM as marked on the scale. n = 145 puncta from 10 chondrocytes in 3 animals. (G-H) Erc1deficient chondrocytes accumulate secretory vesicles at the cell Cortex. Electron micrographs (G) of 3 dpf (EP) and 3.5 dpf (IGP) were used for quantification of cortical vesicle accumulation (H). The boxed region of the second panel is shown in higher magnification in the third panel, and arrowheads point to accumulating vesicles at the cell cortex. n values are number of cells analyzed from at least 2 animals. (I) IF of type II collagen (green) and proteoglycans (WGA, magenta) on EP WT and *kimble* chondrocytes. Arrowheads indicate intracellular localization of ECM cargo in *kimble* chondrocytes. **p<0.01

The cortical localization of Erc1b suggested the possibility that it could regulate vesicle trafficking in chondrocytes. Because chondrocyte cell shape changes coincide with rapid ECM secretion and thus heavy biosynthetic traffic (Fig. 3A-F), we reasoned that membrane and protein cargo delivery to the cell surface could be a potential mechanism for regulating chondrocyte cell shape changes. Thus, we hypothesized that loss of Erc1b impairs exocytic trafficking in chondrocytes. To explore this possibility, we used electron microscopy to determine whether *kimble* chondrocytes accumulate secretory vesicles at the cell cortex. WT chondrocytes displayed abundant organelles indicative of high biosynthetic activity, such as rough endoplasmic reticula (rER) and mitochondria (Fig. 6G). At 3 dpf (EP), the onset of cell shape changes, *kimble* chondrocytes were similar in appearance to wild types, with abundant secretory organelles (Fig. 6G); however, they also contained abundant electron dense vesicles that were concentrated at the cell cortex, (xlJ = 0.597 ± 0.084 µm from the PM, 95% CI, n=121). Cortical vesicles were also present in wild types, but they were > 3-fold more abundant in *kimble* chondrocytes (Fig. 6G,H). Consistent with impaired exocytic trafficking, *kimble* chondrocytes accumulated type II collagen and proteoglycans in the cytoplasm and near the cell cortex (Fig. 6I). Notably, in *kimble* mutants, type II collagen and proteoglycans were also present in the extracellular space and not completely retained intracellularly (Fig. 6I), indicating that while vesicle trafficking is impaired, cargo secretion is not completely abolished in mutants.

If impaired vesicle trafficking in the exocytic pathway is responsible for cell shape dysregulation in *kimble* mutants, then inhibiting components that function in Erc1-dependent exocytosis in WT chondrocytes should phenocopy *kimble* cell shape defects. To test this hypothesis, we mosaically expressed transgenes encoding dominant negative (DN) alleles of Rab8 and Kinesin-1, which both function in exocytic vesicle traffic to Erc1 cortical platforms ^29,30^. We generated mosaic transgenic WT embryos expressing WT or DN *rab8a* or *kif5ba* in isolated clones of transgenic chondrocytes using the *col2*α*1* promoter as a driver (Fig. 7A), and then compared their cell shape to neighboring non-transgenic WT chondrocytes. Strikingly, inhibiting Rab8 or Kinesin-1 function using this strategy recapitulated *kimble* cell shape defects, resulting in constricted chondrocytes immediately adjacent to properly-shaped controls in WT cartilage (Fig. 7B-D). Moreover, as with *kimble* chondrocytes, these cell shape defects were suppressed by partially destabilizing microtubules (Fig. 7B-D). Together, our data suggest that cell shape and cartilage morphology defects in *kimble* mutants stem from dysregulation of trafficking in the exocytic pathway.

**Figure 7.**
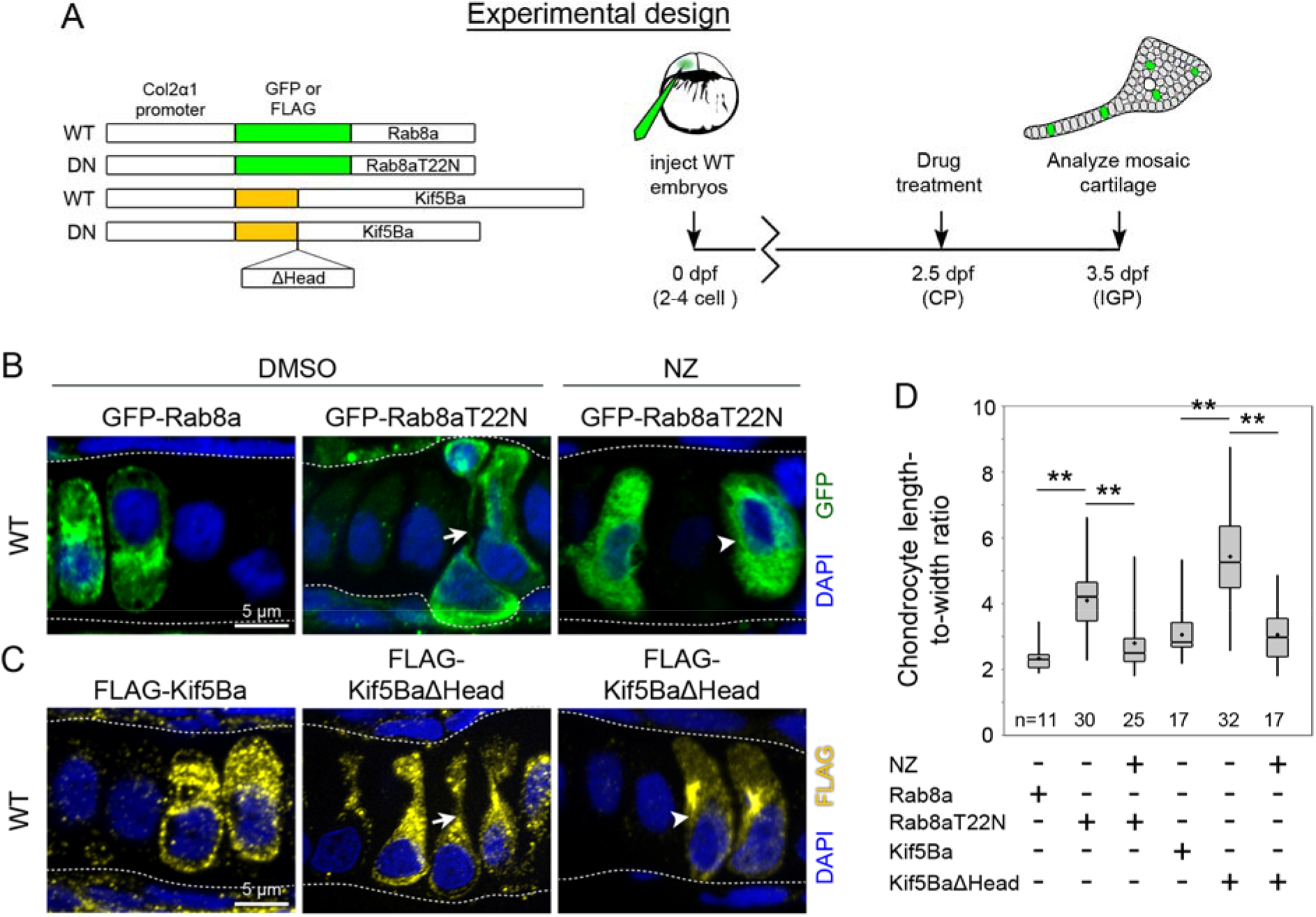
Inhibiting cortical trafficking components independently of Erc1b in WT chondrocytes phenocopies *kimble* cell shape defects. (A) Experimental strategy for mosaically inhibiting exocytic trafficking factors in WT chondrocytes. Transgenes encoding WT eGFP-Rab8a, dominant negative (DN) eGFP-Rab8aT22N, WT FLAG-Kif5Ba or DN FLAG-Kif5BaΔHead were injected into a single blastomere of 2-4 stage embryos. CP-stage embryos were then treated with DMSO (control) or 0.6 µM nocodazole (NZ) for 24 hr before analysis. (B,C) Mosaic expression of Rab8 and Kinesin-1 constructs in WT cartilage. Arrows indicate chondrocytes expressing DN Rab8 and Kinesin-1 that exhibit cell constriction, and arrowheads point to transgenic chondrocytes treated with nocodazole with rescued cell shape. n-values are total number of cells analyzed from a minimum of 4 animals. (D) Quantification of chondrocyte cell shape changes after indicated treatment. **p<0.01

### Imbalance of endo/exocytic recycling drives cell shape defects in the absence of Erc1b

Our genetic data indicate that dysregulation of exocytic vesicle trafficking leads to chondrocyte cell shape defects in *kimble* mutants. However, *kimble* chondrocytes appear to decrease in cell width over time, suggesting that exocytic defects (impaired cargo delivery to the PM) alone cannot explain this phenotype. Because the lateral membrane constricts in mutants, we asked whether this cell shape change is caused by an imbalance of endocytic and exocytic trafficking. To address this question, we first analyzed PM dynamics during chondrocyte cell growth *in vivo* using live imaging to precisely determine the nature of cell shape changes. We crossed the *kimble* line to the *Tg*(*col2*α*1:caax-eGFP)* background, which labels the chondrocyte PM and cytoplasmic structures derived from the PM. We then imaged the medial SY cartilage in 3 dpf (EP) live embryos for > 3 hours to track chondrocyte cell shape changes and PM dynamics (Fig. 8A). WT chondrocytes exhibited subtle PM changes that led to proportional length and width growth over the time course (Fig. 8B-E). Moreover, we also observed many dynamic punctate structures that were prominent at the middle of the cell cytoplasm (Supp. Video 6), suggesting that these structures may derive from the lateral PM. In contrast to wild types, *kimble* chondrocytes decreased in cell width while elongating at a rate comparable to WT cells (Fig. 8B-E). Most *kimble* chondrocytes progressively decreased in width while others exhibited oscillatory patterns of expansion and contraction before ultimately remaining constricted in width (Supp. Video 7). Moreover, as with wild types, *kimble* chondrocytes showed dynamic caax-eGFP+ structures at the middle of the cell, concentrated near the constricting lateral PM (Supp. Video 7).

**Figure 8.**
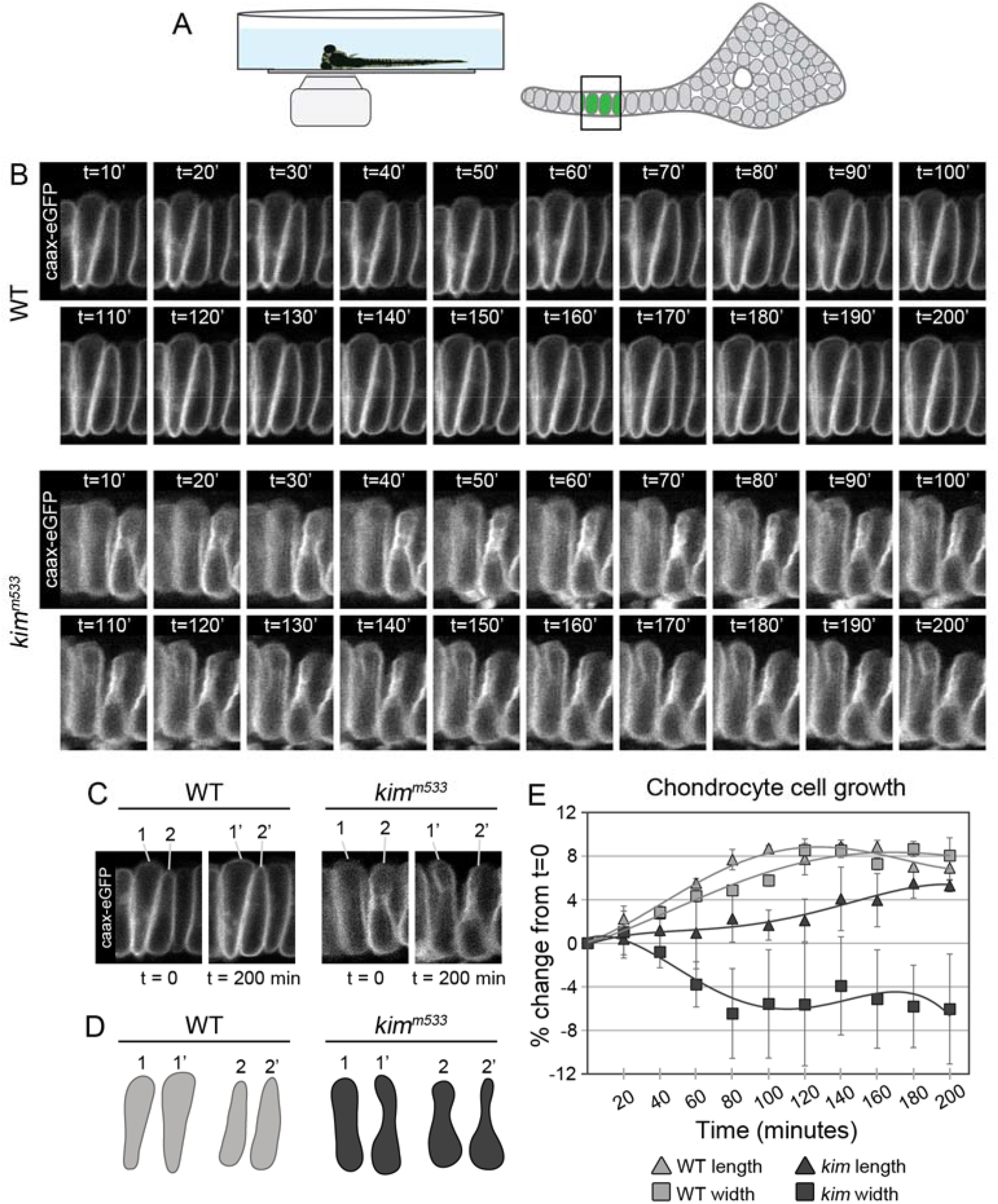
Erc1b is required for isometric plasma membrane growth during cartilage morphogenesis. (A) Experimental strategy for imaging cell shape changes in vivo. Live EP stage (3 dpf) *Tg(col2a1:caax-eGFP)* embryos were immobilized in low melting point agarose and the medial symplectic cartilage was imaged at high magnification by time-lapse confocal microscopy. (B) Frames of time-lapse imaging of chondrocyte cell shape at 10 minute intervals. (C,D) First and last time intervals of live imaging and cell outline tracing to illustrate cell shape changes. (E) Quantification of chondrocyte cell shape changes from time-lapse analysis. WT chondrocytes undergo approximately uniform growth in length and width while *kimble* chondrocytes elongate similar to wild types but laterally constrict in width. n ≥ 6 chondrocytes from at least 2 animals for each genotype.

We hypothesized that during cell growth, endocytic and exocytic recycling impacts cell shape changes, and that *kimble* chondrocytes dysregulate cell shape by altering the balance of activities in this pathway. To test this hypothesis, we attempted to manipulate the recycling pathway in *kimble* chondrocytes by inhibiting the function of Rab11, a well-established factor regulating endocytic recycling in epithelial cells^31^. We generated mosaic transgenic WT or *kimble* chondrocytes expressing either eGFP-Rab11a or DN eGFP-Rab11aS25N. We used this approach to eliminate non-cell-autonomous effects on cell shape and to use neighboring non-transgenic chondrocytes as internal controls for quantitative cell shape analysis. Strikingly, inhibiting Rab11 function suppressed cell shape defects in *kimble* chondrocytes (Fig. 9A,B), and we observed elliptical shaped DN Rab11-expressing *kimble* chondrocytes immediately adjacent to constricted non-transgenic controls (Fig. 9A). Additionally, we found that eGFP-Rab11a was present at cortical puncta and enriched near the lateral PM, marked with WGA (Fig. 9A), suggesting they may represent cortical recycling endosomes. Consistent with this notion, we observed in TEM goblet-shaped invaginations resembling endocytic pits at the lateral PM in 3 and 3.5 dpf (EP, IGP) chondrocytes (Fig. 9C). In summary, we propose that Erc1b regulates endo/exocytic recycling to modulate PM dynamics and cell shape during a heavy phase of biosynthetic traffic and ECM secretion after chondrocyte differentiation (Fig. 9D).

**Figure 9.**
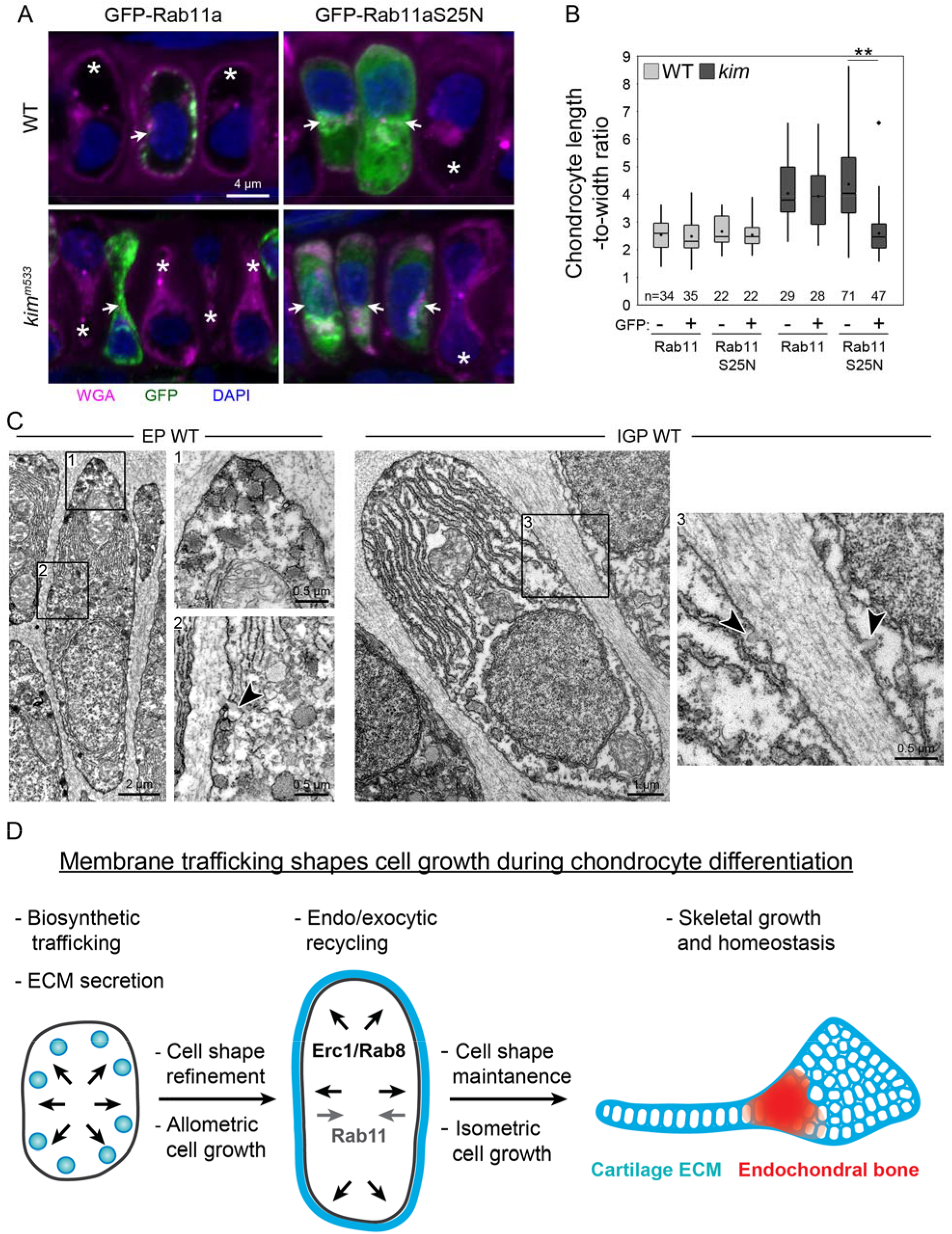
Rab11-dependent recycling pathway mediates chondrocyte cell constriction during impaired exocytosis. (A) Inhibition of Rab11-dependent recycling pathway rescues *kimble* chondrocyte shape cell-autonomously. IF of transgenic (GFP+, white arrows) WT (eGFP-Rab11a) and dominant negative (eGFP-Rab11aS25N), and non-transgenic (GFP-, asterisks) neighboring control chondrocytes. eGFP-Rab11a is enriched at cortical puncta at the lateral membrane. (B) Quantitative analysis of cell shape. n-values are total number of cells analyzed from at least 4 animals. (C) Electron micrographs of 3 dpf (EP) and 3.5 dpf (IGP) WT chondrocytes. EP chondrocytes exhibit electron dense secretory vesicles at polarized ends (inset 1) and the lateral cortex. EP and IGP chondrocytes show goblet-shape endocytic pits at the lateral plasma membrane (insets 2 and 3, arrowheads). (D) Model of Erc1b function in skeletal development and homeostasis. Small round progenitor cells differentiate into highly secretory chondrocytes, delivering biosynthetic cargo (ECM and membrane) to the plasma membrane, and driving cell growth. Regulation of membrane recycling (uptake and delivery) at the cell surface via endo/exocytic trafficking shapes cell growth to promote the highly conserved “stack of coins” chondrocyte cell shape. Dysregulation of this pathway leads to chondrocyte cell death, cartilage dysmorphology and degradation, and impaired endochondral bone formation. **p<0.01, ♦outlier.

## Discussion

Here, we uncovered a developmental mechanism for regulating cell shape changes as chondrocytes elongate after convergent extension to promote cartilage growth. NCC-to-chondrocyte cell differentiation is associated with cartilage ECM secretion and dramatic cell growth, which promotes cartilage growth and craniofacial morphogenesis. Rapid biosynthetic membrane traffic to the plasma membrane and ECM secretion possibly drives chondrocyte cell growth at this phase since mutants that block biosynthetic traffic halts chondrocyte elongation^9,10,32^. Endo/exocytic recycling, on the other hand, shapes and resolves cell growth to promote the conserved “stack-of-coins” architecture that is characteristic of mature chondrocytes. We propose that upon loss of the cortical scaffold protein Erc1b or Rab8 inhibition, imbalance of Rab11-dependent endocytic recycling drives cellular constriction along the mediolateral cell axis. Although chondrocyte intercalation is unaffected, cell shape dysregulation severely impairs cartilage growth and later induces chondrocyte cell death, further altering craniofacial morphogenesis. Our study therefore reveals that cell shape regulation in newly differentiated chondrocytes is a critical, narrow developmental window for ensuring proper growth and morphogenesis of the embryonic head.

Craniofacial malformations are present in approximately one third of all human congenital syndromes^1^, but these disorders are often associated with multiple susceptible loci^3^ or severe cytogenic defects^4^, and their etiologies are not well understood. Given that craniofacial morphogenesis is a complex trait influenced by multiple genes, gene-environment interactions, and gene regulatory landscapes^33^, deciphering the molecular basis of craniofacial malformations is challenging. Therefore, as with other complex disorders^34,35^, animal model studies of gene function during embryonic development are needed to identify causal pathogenic variants underlying human congenital diseases^10,36^. The human *ERC1* gene is located on chromosome 12p13.33. Deletions at this locus are associated with behavioral and learning difficulties^37–39^, orofacial praxia^40^, and craniofacial dysmorphologies^4,37–41^, with varying severity and variable expressivity of inherited alleles among affected individuals and probands. Notably, craniofacial defects are a common phenotype associated with 12p13.33 deletions, and nearly half of publicly available incidents of copy number variants of *ERC1* report similar morphological abnormalities (http://decipher.sanger.ac.uk). However, *ERC1* has not been considered as a candidate disease gene for craniofacial syndromes. Modern genomic approaches are now being applied to understand the genetic basis of craniofacial syndromes^3^. On the basis of our developmental studies and reported cases of 12p13.33 microdeletions, we propose that *ERC1* be prioritized as a candidate genetic factor contributing to craniofacial disorders.

## Materials and methods

### Fish maintenance and breeding

Zebrafish were raised and kept under standard laboratory conditions at 28.5°C as previously described ^42^. All experiments were conducted in accordance with the guidelines established by the IACUC at Vanderbilt University Medical Center.

### Genetic mapping and cloning

The *kimble* locus was mapped in an F2 intercross using bulked segregate analysis. DNA samples were PCR-genotyped with SSLP markers evenly spaced across the zebrafish genome ^21^. The mapped *kim*^*m533*^ mutation was confirmed by Sanger sequencing of genomic DNA flanking the mutation site from two homozygous wild-type F2 animals, six heterozygous F2 animals, six homozygous mutant F2 animals and two animals each from three different genetic backgrounds of wild-type fish (AB, IN and TL) ^43^.

### Cartilage proteoglycan staining

Alcian Blue staining was performed as previously described ^32^ with modifications. Larvae were fixed in 4% PFA, stained overnight in 0.1% Alcian Blue in 70% EtOH/1% HCL, de-stained in 70% ethanol/ 5% HCl, and then cleared in 50% glycerol/0.25% KOH. Differential interference contrast (DIC) images were acquired from dissected, flat-mounted cartilage using a 100x oil-immersion objective mounted on a Zeiss AxioImager Z1.

### Morphometric analysis of chondrocyte cell shape

Average cell width was calculated by measuring cell length and area in ImageJ and using the formula for the area of an ellipse, A=π*a*b, where a is the length of the major axis (cell length/2) and b is the length of the minor axis (cell width/2). Thus, average cell width was calculated using the formula, width=(4*area)/(π*length).

### Cell death analysis

Double strand DNA breaks were detected using the TUNEL method as previously described ^44^. Embryos were anesthetized, fixed in 4% PFA, and then processed for cryosectioning. TUNEL was performed using the Roche *In Situ* Death Detection Kit.

### Electron microscopy

Samples were processed as previously described ^11^. Embryos were fixed in 2.5% glutaraldehyde in 0.1 M sodium cacodylate at room temperature for 1 hr. and then overnight at 4°C. After rinsing in 0.1 M sodium cacodylate, samples were post-fixed with 1% osmium tetroxide in 0.1 M sodium cacodylate for 1 hr. Following additional rinsing, specimens were dehydrated step-wise in ethanol and then propylene oxide, infiltrated with resin step-wise, and then embedded in resin for 48 hours at 60°C. 50 nm sections were collected on a Leica Ultracut Microtome and analyzed on a Phillips CM-12 Transmission Electron Microscope provided by the VUMC Cell Imaging Shared Resource.

### Immunofluorescence (IF), fluorescence microscopy, and image processing

Processing for IF was performed as previously described ^10^ with modifications. Embryos were fixed in freshly prepared 2-4% PFA and incubated in 30% sucrose/PBS prior to embedding in Cryomatrix, (Thermo Scientific). Frozen sections were collected with a Leica CM-1900 cryotome. Primary antibodies used were GFP (Vanderbilt Antibody and Protein Resource), Erc1 (Abcam), alpha-tubulin (Abcam), and FLAG, M2 clone (Sigma). Alexa Fluor 488-and 555-(Life Technologies) or DyLight 550-conjugates (Thermo Scientific) were used for secondary antibodies. To-Pro-3 (Molecular Probes) or 4’,6-diamidino-2-phenylindole (DAPI, Molecular Probes) was applied as a nuclear counterstain. Proteoglycan and cortical staining was performed using Alexa Fluor-conjugated Wheat Germ Agglutinin (WGA, Life Technologies) and Phalloidin (Life Technologies), respectively. Slides were mounted in ProLong Gold (Life Technologies) prior to imaging, which was performed using Zeiss LSM-510 Meta or LSM-710 Inverted Confocal Microscopes (VUMC Cell Imaging Shared Resource) with a Plan-Apochromat 63x/1.40 Oil objective or an AxioImager Z1 equipped with an Apotome (Zeiss) and an EC Plan Neofluar 100x/1.30 Oil objective. Widefield fluorescence data were deconvolved using default settings using the Zen 2012 software package (Zeiss). For tubulin IF, samples were fixed in a buffer containing PFA and glutaraldehyde in Cytoskeletal Buffer (9 mM MES (2-(4-Morpholino)ethane Sulfonic Acid), 150 mM NaCl, 5 mM EGTA, 5 mM MgCl_2_, 5 mM glucose), quenched in 0.1% NaBH_4_, and then processed as described above. Post-acquisition data processing was limited to linear adjustments for brightness using the levels tool in Adobe Photoshop.

### Distance measurements of Erc1b puncta from the cell membrane

Embryos were fixed for 20 minutes in 2% PFA and then processed for cryosectioning, IF, and confocal microscopy as described above. Z-stacks were processed in Imaris 8.0 (Bitplane) with standard Matlab extensions. The caax-eGFP channel was used to create a surface of the cell membrane of individual chondrocytes, and the *spots* tool was used to detect Erc1b puncta. The distance transformation extension was used to measure the distance of Erc1b puncta from the cell membrane surface and to pseudo-color Erc1b spots. Distance measurements of 10 cells from 3 animals were collected using the statistics tab.

### 3D rendering of whole cartilage elements

Embryos were anesthetized and fixed in 4% PFA overnight, bleached in H_2_O_2_ /KOH solution, permeabilized, and then stained with Alexa Fluor 488-conjugated Wheat Germ Agglutinin (WGA, Life Technologies) overnight. Embryos were then rinsed, cleared in glycerol overnight, mounted in agarose on glass bottom dishes, and confocal z-stacks of the entire hyosymplectic (HS) cartilage (approximately 80-100 μm thick) were collected using a Zeiss LSM-710 Inverted Confocal Microscope using a Plan-Apochromat 20x/0.8 M27 objective. 3D projections, rendering, and volume measurements were performed in Imaris 8.0 using the surfaces tool and statistics tab.

### Generation of transgenic constructs

Transgenic constructs were generated using gateway cloning and the Tol2kit ^45^ with primers listed in Supplementary Table 1. pME-eGFP-Rab11a and pME-eGFP-Rab11aS25N, generated by the Link laboratory ^46^ and shared with us by the Bagnat laboratory ^47^, were used for assembly of our Rab11 constructs.

### Whole mount *in situ* hybridization

Whole mount *in situ* hybridization with probes recognizing *sox9a* and *col2*α*1* was performed as previously described ^42^. The *erc1b* probe was made by cloning 1041 nucleotides of 3’ UTR from cDNA into the pGEM-T Easy Vector (Promega) using primers listed in Supplementary Table 1.

### Histology

Processing for histology from JB-4 plastic resin was performed as previously described ^9^. Sections were stained with Toluidine blue (Sigma).

### Embryo lysis and western blotting

Epiboly stage embryo lysates were prepared as previously described ^48^ with modifications. Embryos were dechorionated with Pronase (Merck), de-yolked, lysed in RIPA buffer supplemented with protease inhibitor cocktail (RIPA+PI) by rocking at 4°c, and then clarified by centrifugation. For later-stage embryos, whole heads were dissected at the ear capsule, pestle-homogenized in RIPA+PI, and then prepared as described above. Lysates were boiled for 10 min in Laemmli sample buffer, separated by SDS-PAGE, and transferred to PVDF membranes (BioRad). Membranes were blocked with 2% non-fat milk in TBS–Tween (50 mM Tris, pH 7.4, 150 mM NaCl, 0.1% Tween-20), incubated overnight with primary antibody in 2% BSA in TBS-Tween at 4 °C, followed by peroxidase-conjugated secondary antibodies (Promega). Signal was developed using ECL substrate (Perkin Elmer), and detected using a ChemiDoc system (BioRad). Antibodies used were Erc1 (Abcam) and alpha tubulin, DM1A clone (Sigma).

### Live imaging of chondrocyte cell shape changes

3 dpf WT or *kim*^*m533*^ *Tg(col2*α*1:caax-eGFP)* embryos were anesthetized in tricaine and mounted in 1.2% low melting point agarose in 0.3x Danieux buffer. Embryos were positioned laterally, and confocal images were collected every 2 minutes for 4 hours using an LSM710 (Zeiss) with a 63x 1.4 NA objective. Cell length and area were measured using ImageJ.

## Statistical analysis

For comparisons of 2 groups, p-values were calculated using two-tailed unpaired t-tests. For comparisons of more than 2 groups, one-or two-way analysis of variance (ANOVA) was used to test for significance followed by Tukey’s test to determine p-values. Categorical data was analyzed using Fisher’s exact test. All bar graphs are presented as mean ± SD.

## Acknowledgements

We thank Cory Guthrie for excellent animal care, Witold Rybski and Kirill Zavalin for technical assistance, Xiaodong Zhu, Jacek Topczewski, Mike Dohn, Jason Jessen, Michel Bagnat, and Josh Gamse for sharing reagents, and Wolfgang Driever for the kimble mutant line. Thanks to Anne Kenworthy, Florent Elefteriou, Todd Graham, Irina Kaverina, David Miller, and Wendell Yarbrough for helpful advice. The Vanderbilt University Cell Imaging Shared Resource (CISR) provided electron microscopy services. This work was supported in part by the Zebrafish Initiative of the Vanderbilt University Academic Venture Capital Fund, the NIH NIDCR grant R01 DE018477 (E.W.K.). D.S.L. was supported by NRSA F31DE022226 and T32HD007502 Training Program in Developmental Biology; D.B.M was supported by T32GM008554, the Cellular, Biochemical and Molecular Sciences Training Program; G.U. was supported by the Vanderbilt International Scholar Program, and American Heart Association predoctoral fellowship 15PRE22940041. The Vanderbilt University Cell Imaging Shared Resource is supported by NIH grant DK020593

## Author contributions

Author contributions

Conceptualization: D.S.L., E.W.K.; Methodology: D.S.L., G.U., D.B.M.; Validation: D.S.L.; Formal analysis: D.S.L., G.U.; Investigation: D.S.L.; Resources: E.W.K.; Writing - original draft: D.S.L., E.W.K.; Writing - review & editing: D.S.L., E.W.K.; Visualization: D.S.L.; Supervision: E.W.K.; Project administration: E.W.K.; Funding acquisition: E.W.K.

---

**Supplementary Figure 1.**
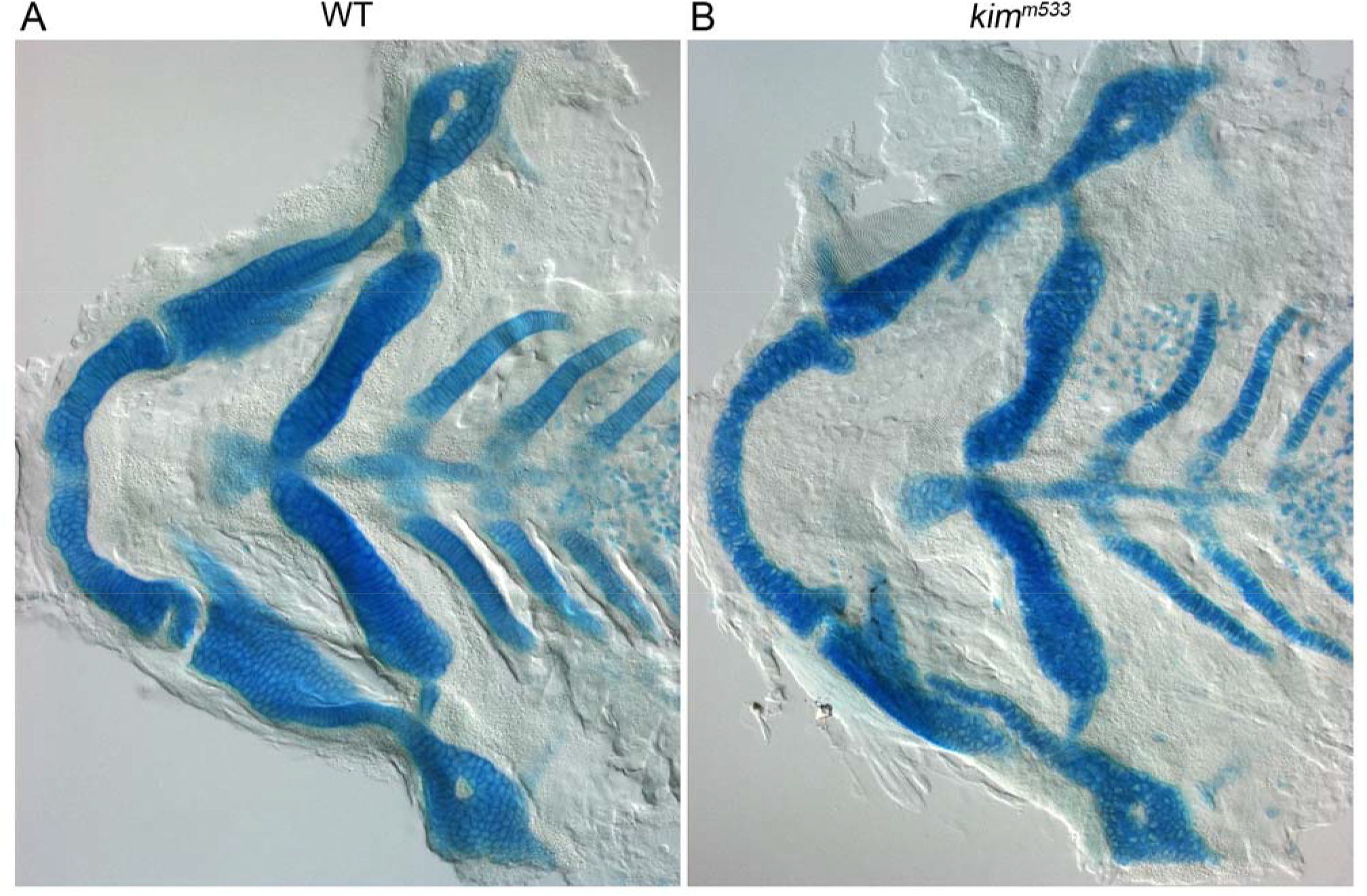
Cartilage morphology in embryonic-stage *kimble* mutants. (A,B) DIC imaging of flat-mounted Alcian-blue stained skeletal preparations of 3.5 dpf WT (A) and *kimble* mutant embryos (B). Gross defects in skeletal patterning are not obvious at this stage in mutants.

**Supplementary Figure 2.**
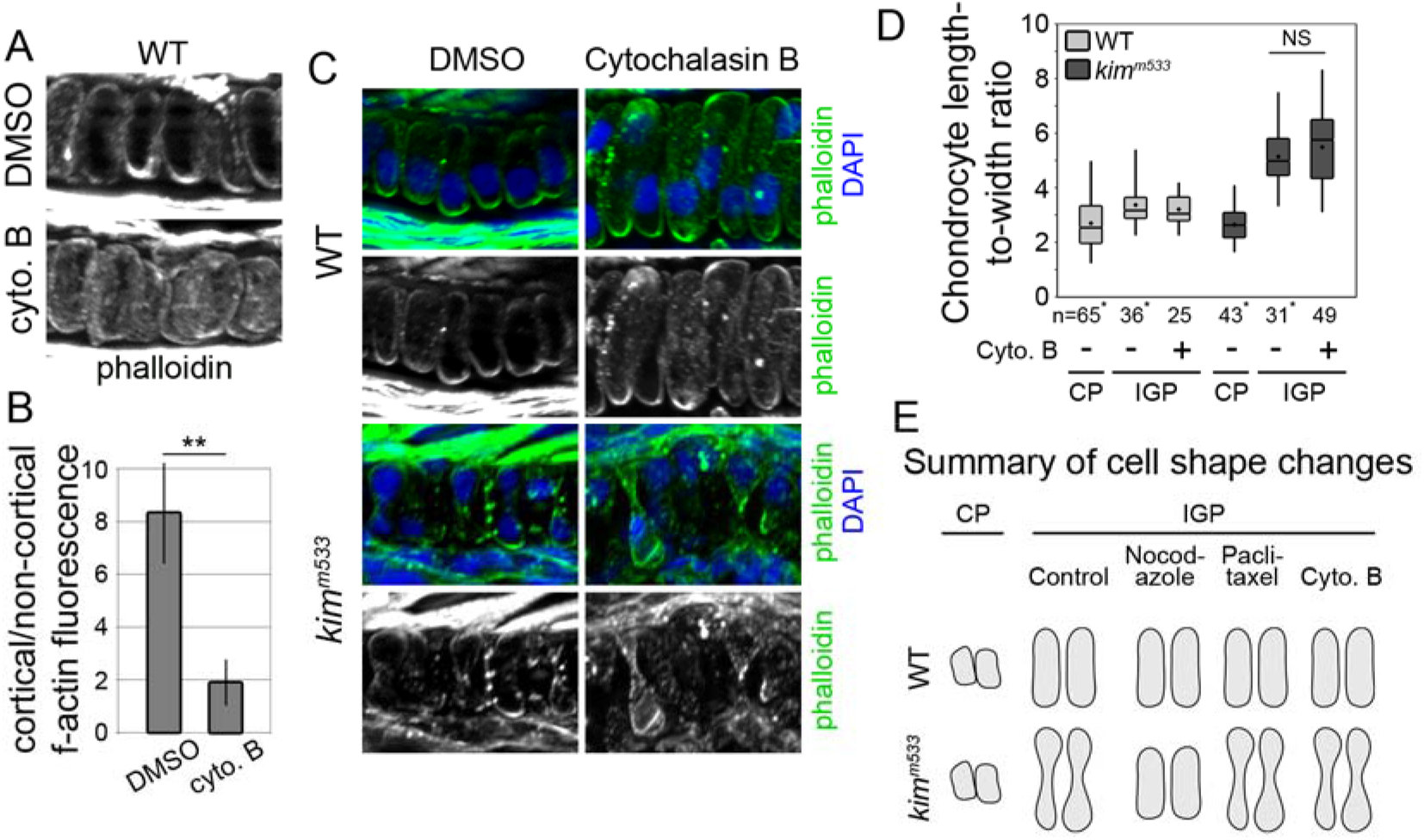
Cortical F-actin disruption does not impact *kimble* chondrocyte cell changes. (A) Phalloidin staining of F-actin in chondrocytes after CP-IGP treatment with 1% DMSO or 4 µM cytochalasin B (cyto. B). Cytochalasin B treatment led to increased non-cortical labeling with phalloidin (F-actin redistribution) compared to DMSO-treated chondrocytes. (B) Quantification of phalloidin cortical enrichment. Data are an average ratio of cortical (0-500 nm from cell edge) fluorescence intensity relative to non-cortical, cytoplasmic (> 500 nm from cell edge) intensity. (C) Cortical (phalloidin) and nuclear (DAPI) labeling of WT and *kim*^*m533*^ embryos after CP-IGP treatment with 1% DMSO or 4 µM cytochalasin B. (D) Quantification of chondrocyte cell shape after 4 µM cytochalasin B treatment. Quantitative data for control groups (non-cytochalasin B treated) are the same data set as presented in Fig. 5D. (F) Schematic summary of cell shape changes after cytoskeletal inhibitor treatment. **p<0.01.

**Supplementary Figure 3.**
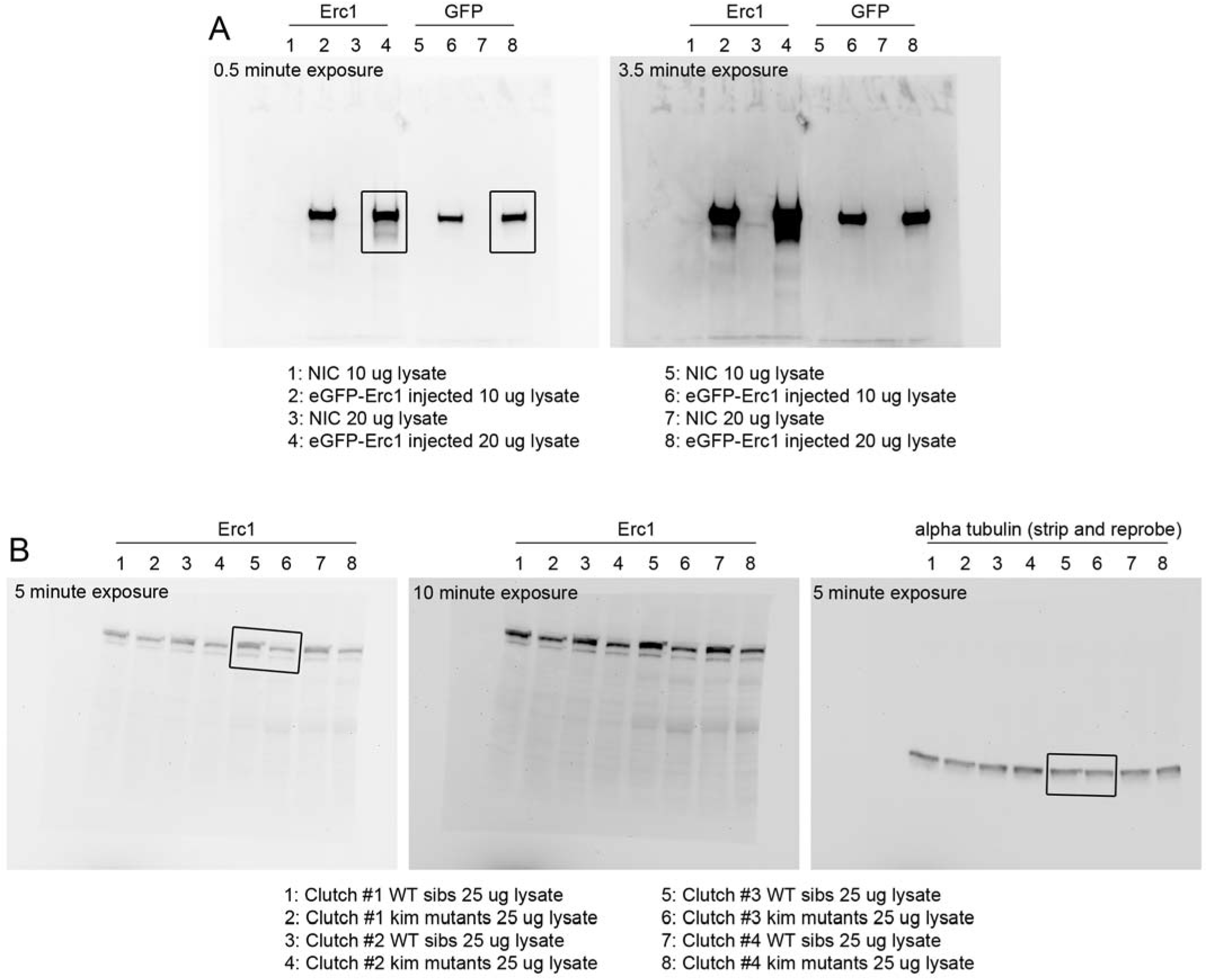
Unedited western blot images. (A) Western blot data from Fig. 6A. (B) Western blot data from Fig. 6B. Boxes represent cropped areas shown in Fig. 6.

**Supplementary table 1.**
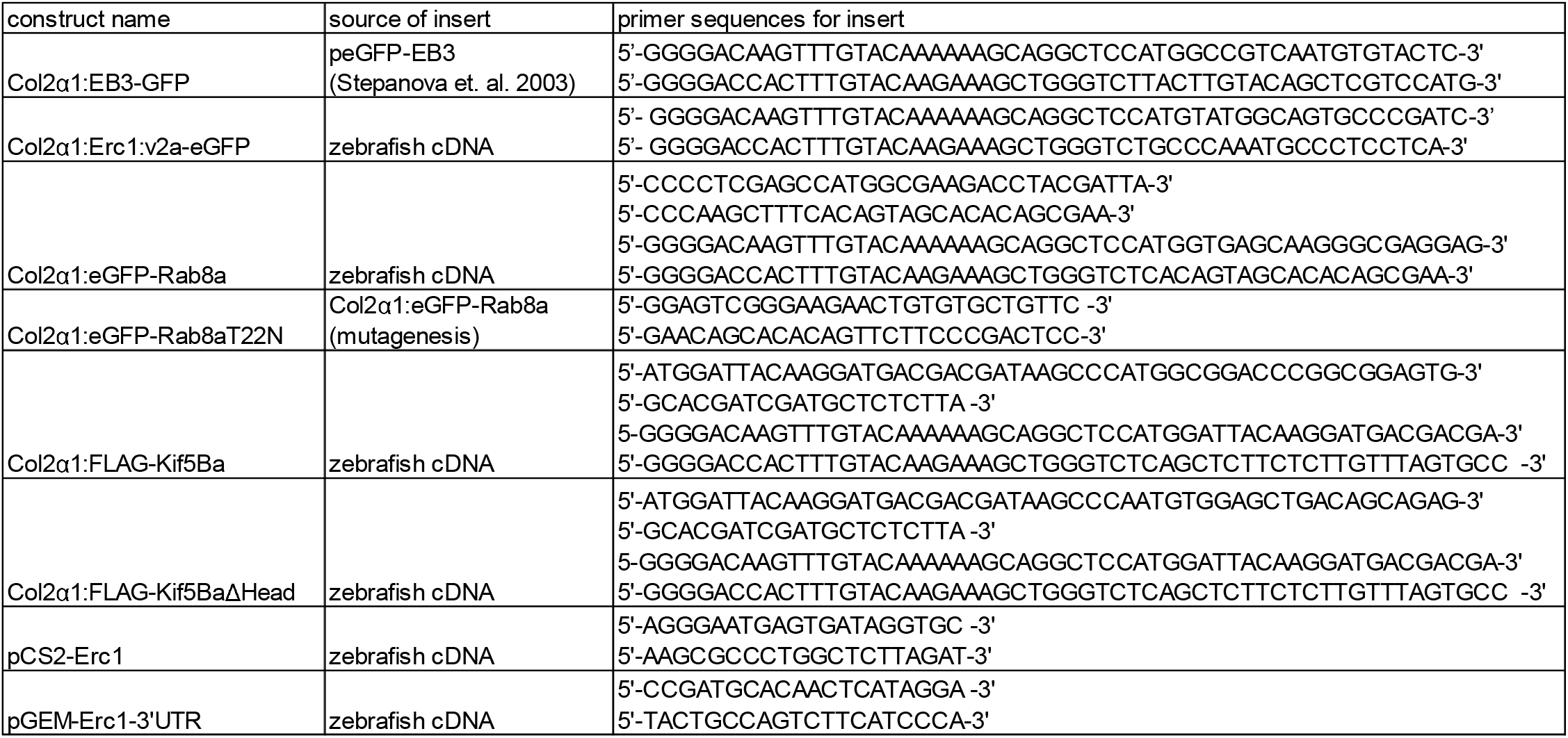
Primer sequences used for cDNA cloning.

**Supplementary Video 1. WT chondrocyte 3D cell shape**

Rotating view of WT chondrocytes from Fig. 4B.

**Supplementary Video 2. *Kimble* chondrocyte 3D cell shape**

Rotating view of *kim*^*m533*^ chondrocytes from Fig. 4B.

**Supplementary Video 3. WT hyosymplectic cartilage 3D shape**

Rotating view of WT HS cartilage from Fig. 4C.

**Supplementary Video 4. *Kimble* hyosymplectic cartilage 3D shape**

Rotating view of *kim*^*m533*^ HS cartilage from Fig. 4D.

**Supplementary Video 5. Erc1b localization in WT chondrocytes**

Rotating view of Erc1b distance mapped Erc1b voxels in WT chondrocytes from Fig. 5J.

**Supplementary Video 6. Live imaging of WT chondrocyte cell growth**

Video of 3 dpf WT chondrocytes from Fig. 8B. Confocal images were collected at 1 frame per minute for 200 minutes.

**Supplementary Video 7. Live imaging of *kimble* chondrocyte cell growth**

Video of 3 dpf *kimble* chondrocytes from Fig. 8B. Confocal images were collected at 1 frame per minute for 200 minutes.

